# A constitutively expressed fluorescence ubiquitin cell cycle indicator (FUCCI) in axolotls for studying tissue regeneration

**DOI:** 10.1101/2021.03.30.437716

**Authors:** Timothy J Duerr, Eun Kyung Jeon, Kaylee M Wells, Antonio Villanueva, Ashley W Seifert, Catherine D McCusker, James R Monaghan

**Affiliations:** Northeastern University, Department of Biology, Boston, MA; University of Massachusetts Boston, Department of Biology, Boston, MA; University of Kentucky, Department of Biology, Lexington, KY

**Keywords:** Regeneration, FUCCI, cell cycle, axolotl

## Abstract

Regulation of cell cycle progression is essential for cell proliferation during regeneration following injury. After appendage amputation, the axolotl (*Ambystoma mexicanum*) regenerates missing structures through an accumulation of proliferating cells known as the blastema. To study cell division during blastema growth, we generated a transgenic line of axolotls that ubiquitously expresses a bicistronic version of the Fluorescent Ubiquitination-based Cell Cycle Indicator (FUCCI). We demonstrate near-ubiquitous expression of FUCCI expression in developing and adult tissues and validate these expression patterns with DNA synthesis and mitosis phase markers. We demonstrate the utility of FUCCI for live and whole-mount imaging, showing the predominantly local contribution of cells during limb and tail regeneration. We also show that spinal cord amputation results in increased proliferation at least 5 mm from the injury. Finally, we use multimodal staining to provide cell type information for cycling cells by combining fluorescence *in-situ* hybridization, EdU click-chemistry, and immunohistochemistry on a single FUCCI tissue section. This new line of animals will be useful for studying cell cycle dynamics using *in-situ* endpoint assays and *in-vivo* imaging in developing and regenerating animals.

**Summary statement:** We generated a ubiquitous transgenic fluorescence ubiquitin cell cycle indicator (FUCCI) axolotl line for examination of cell cycle dynamics during tissue regeneration.

## Introduction

Vertebrate tissue regeneration inherently requires cell proliferation either through endogenous stem cell proliferation or re-entry of differentiated cells into the cell cycle. One of the most striking examples of vertebrate regeneration is epimorphic replacement of the amputated salamander appendage. Appendage regeneration requires the generation of a highly proliferative mass of cells called the blastema. The formation of the blastema is dependent on an intact nerve supply and a specialized layer of epithelium known as the apical epithelial cap (AEC) (McCusker et al., 2015a). The AEC likely has multiple functions including directing outgrowth, maintaining proliferation, and secreting factors that allow for remodeling of underlying extracellular matrix (Stocum, 2017; Tsai et al., 2020). Although the blastema consists of numerous cell types, most cells originate from mesenchymal cell populations located near the amputation plane (Butler, 1933; McHedlishvili et al., 2007). Understanding the mechanisms that initiate and sustain proliferation of blastema cells is a fundamental problem that requires modern molecular tools to track and characterize blastema cell behavior (Stocum, 2017; Tanaka, 2016). Although recent developments in transgenesis and tissue grafting techniques has allowed the observation of blastema cells *in vivo* (Currie et al., 2016; Khattak et al., 2013; Kragl and Tanaka, 2009; Sandoval-Guzmán et al., 2014), further development of transgenic lines are needed to enable imaging of the regeneration process.

In 2008, Sakaue-Sawano and colleagues developed the fluorescent, ubiquitination-based cell cycle indicator (FUCCI) system to study cell cycle progression in human cell lines and mice (Sakaue-Sawano et al., 2008). Since then several variations have been made to the FUCCI construct, and it has been used to generate additional transgenic plants (Yin et al., 2014) and animals (Abe et al., 2013; Sugiyama et al., 2009; Zielke et al., 2014). The FUCCI system is based upon the inverse oscillation of Geminin and Cdt1 proteins that occurs naturally during the cell cycle (Nishitani et al., 2004). The FUCCI construct includes a constitutively active promoter driving expression of a fluorescent protein fused to the Cdt1 protein degron, which has high levels in G1 due to ubiquitin-mediated proteolysis during S, G2 and early mitotic phases (S/G2/M). Conversely, the Geminin protein degron is fused to a different fluorescent molecule that is degraded during late M and G1, leading to high fluorescent protein levels in S/G2/early M ((Zielke and Edgar, 2015) for review). Fusion of these two expression cassettes into a single bicistronic transgene allows visualization of cells while they progress through the cell cycle.

In this study, we use a ubiquitously expressed bicistronic FUCCI construct to generate a transgenic line of axolotl salamanders to study cell cycle dynamics during development, limb regeneration, and tail regeneration.

## Results

### Generation and characterization of developing FUCCI axolotls

We chose to design a bicistronic version of the original FUCCI construct because it should theoretically lead to equimolar levels of probes and only require the generation of a single transgenic animal line (Rajan et al., 2018). The construct includes a CAG promoter that drives expression of monomeric azami green fused to the zebrafish geminin degron (mAG-zGem) followed by a viral 2A self-cleaving peptide and mCherry fused to the zebrafish Cdt1 degron (mCherry-zCdt1), which was cloned into the pISce-Dest backbone (gift of Jochen Wittbrodt) using Gateway cloning (Fig. 1A). F0 animals were generated using standard axolotl injection conditions with I-SceI Meganuclease (Khattak et al., 2009), generating F0 animals with mosaic FUCCI expression. A single female was selected due to strong ubiquitous FUCCI expression (Fig. 1B) to mate with a d/d white male, which generated clutches consisting of 82% and 92.7% transmission rate suggesting the FUCCI construct integrated at multiple sites in the founder animal.

**Figure 1:**
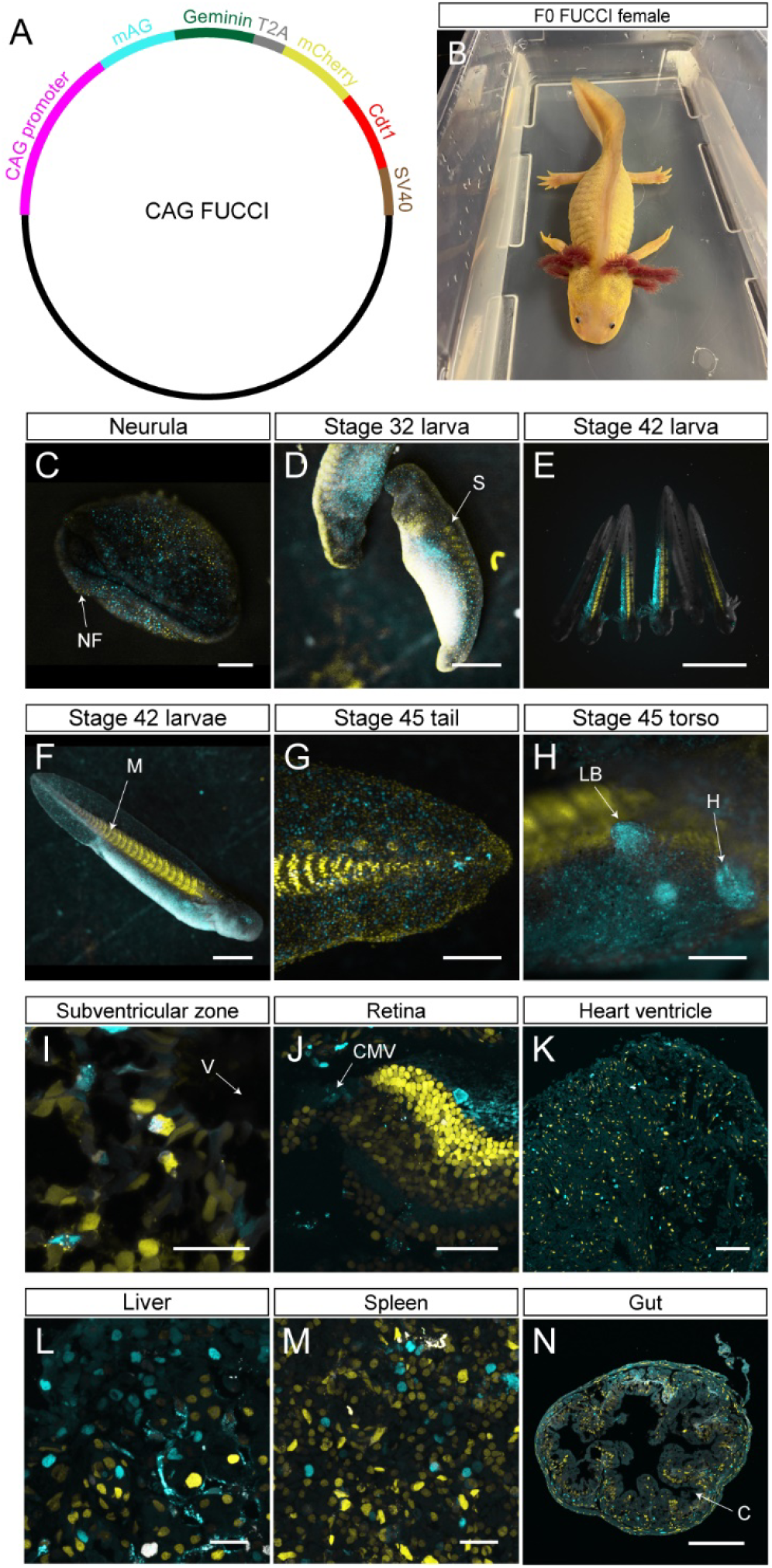
FUCCI probes are expressed in developing and adult, homeostatic tissue. (A) Plasmid map for CAG FUCCI. (B) Sexually mature, F0 FUCCI female crossed with white d/d males to generate the F1 clutch used in the study. (C) Stage 17 neurula expressing FUCCI probes. NF= neural fold. Scale bar= 500 μm. (D) Stage 32 larva. S= somite. Scale bar= 1 mm. (E) Six stage 42 larvae with negative, ubiquitous, and variable expression patterns. Scale bar= 5 mm. (F) Individual stage 42 larva. M= myomeres. Scale bar= 1 mm. (G) Posterior tail tip of a stage 45 larva. Scale bar= 500 μm. (H) Torso of a stage 45 larva. LB= limb bud. H= heart. Scale bar= 500 μm. (I) Subventricular zone of the adult brain. V= ventricle. Scale bar= 50 μm. (J) Adult retina. CMZ= ciliary marginal zone. Scale bar= 100 μm. (K) Adult heart ventricle. Scale bar= 100 μm. (L) Adult liver. Scale bar= 50 μm. (M) Adult spleen. Scale bar= 50 μm. (N) Adult gut. C= crypt. Scale bar= 200 μm. Individual channels for I-J are available with EdU staining in Fig. S1.

Examining live transgenic embryos, we first detected FUCCI protein expression at neurulation with increasing expression throughout development (Fig. 1C-D). Expression was variable between siblings, possibly due to varying levels of transcriptional activation or due to multiple integrations of the FUCCI construct into the genome (Fig. 1D-F). Gross observation of transgenic larvae clearly showed distinct non-proliferative G1 populations including the somites (Fig. 1D-F), the tail myotomes (Fig. 1G), lateral line neuromasts (Fig. 1G), and highly proliferative S/G2/M populations such as the limb bud and larval heart (Fig. 1H). To determine the adult expression pattern of CAG-FUCCI, tissues sections were analyzed from the brain, eye, heart, liver, spleen, and gut. mAG and mCherry expression were observed in every tissue type with little overlap between probes except for differentiated muscle fibers (Fig. 1I-N, Fig. S1).

### FUCCI cell cycling probes overlap with S and M phase markers

To determine the overall expression level of FUCCI, we quantified 2547 cells in fixed regenerating FUCCI spinal cords sections 14 days post amputation (dpa) pulsed with 5-ethynyldeoxyuridine (EdU) for three hours (n=9 animals). In total, 22.34% of the cells were mAG^+^/mCherry^−^, 71.22% were mAG^−^/mCherry^+^, 2.75% were mAG^+^/mCherry^+^, and 3.69% mAG^−^/mCherry^−^ (Fig. S2A). We next determined whether mAG^+^ cells were specific to S phase by performing click-it based EdU detection of DNA synthesis on the same regenerating spinal cord tissues (Fig. 2A). Of the 532 EdU^+^ cells (20.9% of total cells), 88.93% were mAG^+^/mCherry^−^, 3.00% were mAG^+^/mCherry^+^, 1.88% were mAG^−^/mCherry^+^, and 6.19% were mAG^−^/mCherry^−^ (Fig. 3B). Conversely, 76.68% of mAG^+^ cells were EdU^+^ and 23.32% of mAG^+^ cells were EdU^−^ (Fig. S2B), suggesting that that the majority of mAG^+^ cells were in S phase rather than G2. This suggests that the S phase is longer than the combined G2/M phase by approximately three fold, which is supported by previous studies (McCullough and Tassava, 1976). In order to study mitosis in a highly proliferative tissue, we performed immunohistochemistry for phosphorylated serine 10 histone H3 (pHH3) in 10 dpa regenerating limb blastemas (n=3) (Fig S2C-F). We found that 89.13% percent of pHH3^+^ cells were mAG^+^/mCherry^−^, 2.17% were mAG^+^/mCherry^+^, 0% were mAG^−^/mCherry^+^, and 8.69% were mAG^−^/mCherry^−^ (Fig. 2C, Fig S2C-F). As expected, some pHH3^+^ had neither probe signal since Geminin is known to degrade in the late stages of M stage (McGarry and Kirschner, 1998). Overall, the high correspondence between EdU and pHH3 with mAG+ expression shows that mAG-zGem effectively marks cells in both S and M phase.

**Figure 2:**
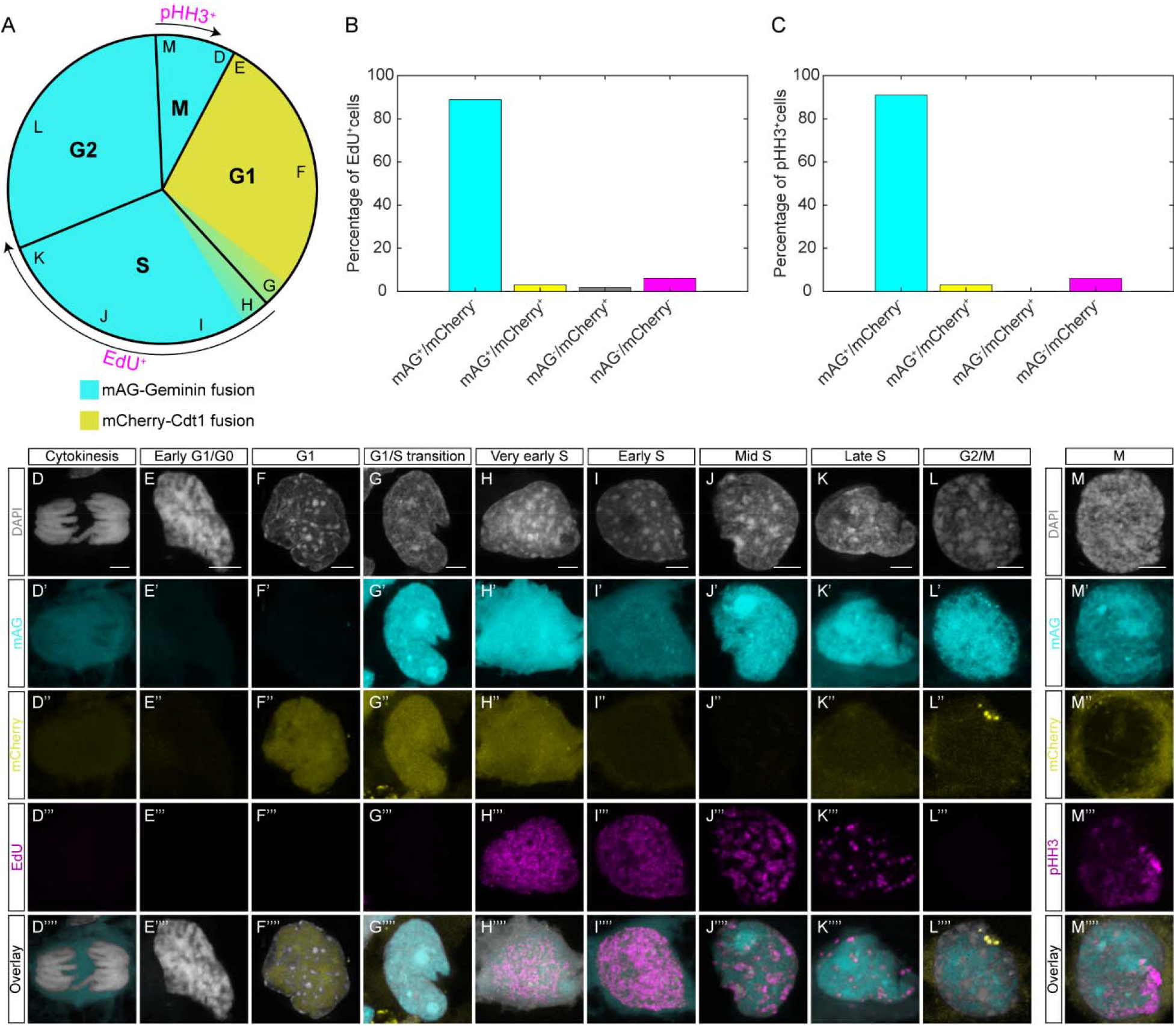
Validation of FUCCI expression with EdU and pHH3. (A) Schematic of the cell cycle with expected staining patterns of EdU and pHH3. Note that EdU may label cells in early G2 as a result of a three hour chase and that pHH3 weakens during late M phase. Letters in the outer edge of the schematic represent the location in the cell cycle of cells from panels D-M. (B) Characterization of EdU^+^ cells in 14 dpa regenerating spinal cords. (C) Characterization of pHH3^+^ cells in 10 dpa regenerating limb blastemas. (D-L) Individual cells from EdU pulsed tissue at every cell cycle stage. Scale bars= 5 μm. (M) Individual cell in M stage from tissue stained for pHH3. Scale bar= 5 μm.

**Figure 3:**
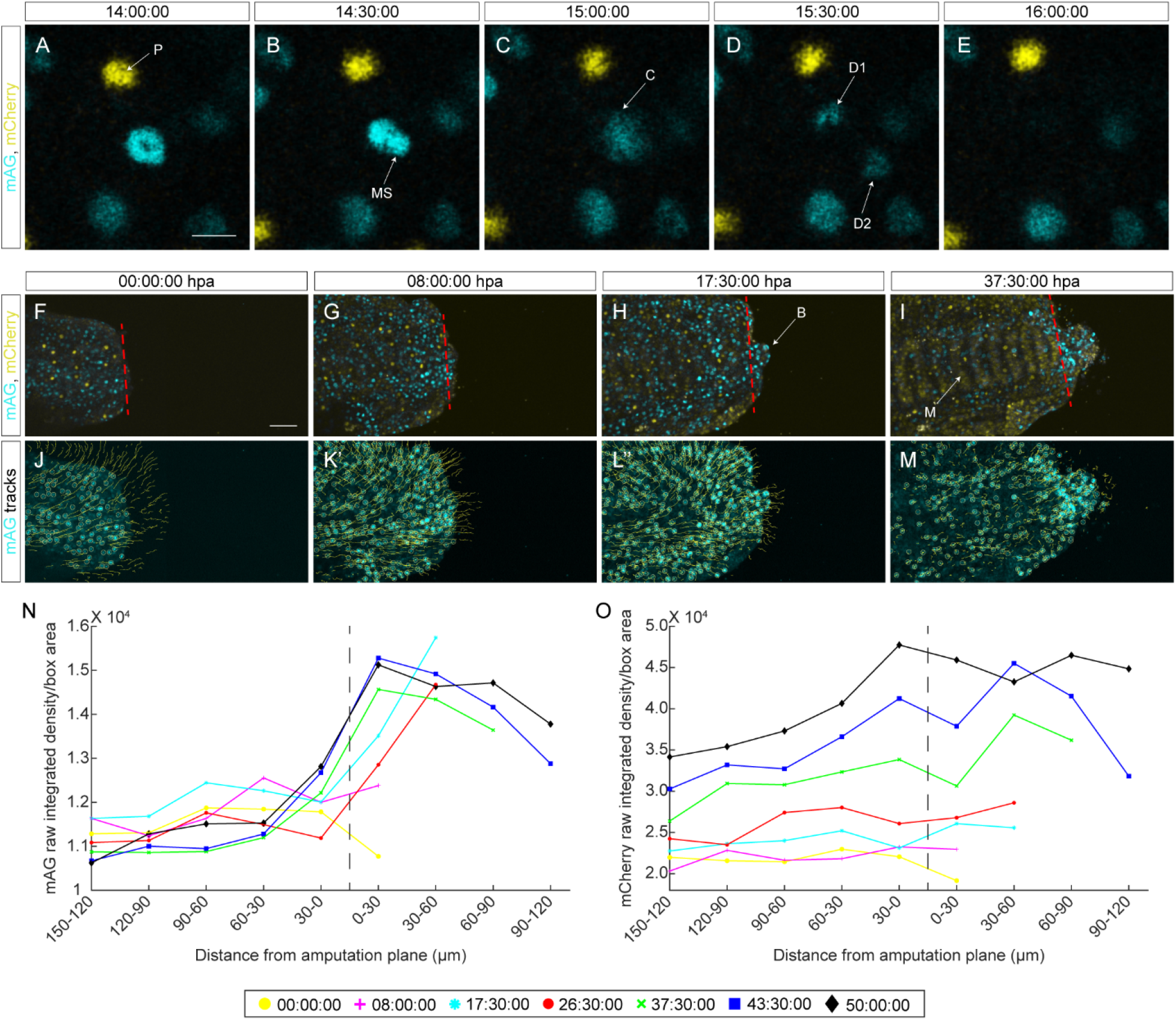
Continuous live imaging of FUCCI tissue. (A-E) Two hour time lapse in five, 30 minute intervals of a diving epithelial cell from a stage 32 larva. P= pigment cell. MS= mitotic spindle. C= cytokinesis. D1= daughter cell 1. D2= daughter cell 2. Scale bar= 25 μm. (F-I) Four frames from the 60 hour live image depicting a regenerating tail after amputation (F), after wound healing (G), during blastema formation (H), and during blastema growth (I). B= blastema. M= myomeres. Scale bar= 50 μm. (J-M) Tracks depicting cell migration in the frames from (J-M). Each line represents the path a cell took 20 frames prior to the current frame and 20 frames after. (N-O) Charts depicting mAG raw integrated density/area (N) or mCherry raw integrated density/area (O) for seven frames from the 60 hour live image. Measurements were obtained by dividing the anteroposterior axis of the regenerating tail into boxes with a width of 30 μm (Fig. S3). The vertical dotted line represents the amputation plane.

We next identified cells at each stage of the cell cycle according to DAPI and EdU staining. Each stage of the cell cycle was observed in proliferating tissues and had predictable genomic structure, EdU incorporation, pHH3 staining, and FUCCI reporter expression (Fig. 2D-M). Collectively, these results demonstrate that our FUCCI construct correctly and reproducibly labels specific stages of the cell cycle.

### Real-time *in-vivo* imaging of FUCCI expression

To determine the feasibility of real-time, *in-vivo* imaging of FUCCI tissue, we imaged cycling epithelial cells in an anesthetized stage 32 larva mounted in 0.3% agarose (Fig. 3A-E, Movie 1). Here we observed that dividing mAG^+^ cells complete the process of mitosis in under 30 minutes to produce two daughter cells with fading mAG intensity (Fig. 3A-E, Movie 2). During this process, we observe both the formation of the mitotic spindle in prophase and cytokinesis after chromosome separation. We did not observe any cell transition from mAG to mCherry or vice versa. This is likely due to the shortening of the G1 phase during embryonic development (Siefert et al., 2015), which would prevent the accumulation of mCherry protein to provide a detectable signal while transitioning from M phase or prior to transitioning to S phase.

To determine the origin of cells during tail regeneration, we performed live imaging of a regenerating tail in a stage 36 FUCCI animal. After amputation, the anesthetized larva was immediately embedded in 0.3% agarose and imaged every 30 minutes over 60 hours (Movie 3). During this 60 hour imaging experiment, we observed early wound healing (Fig. 3G), blastema formation (Fig. 3H), and myomeric muscle development (Fig. 3I). By 8 hours post amputation (hpa), the tail stump was completely covered by a thin layer of epithelium that had both mAG^+^ cells and mCherry^+^ cells (Fig. 3G). Shortly after, an early blastema was observed in the posterior tail tip by 17.5 hpa (Fig. 3H), composed mostly of mAG^+^ cells. At this time point, early myomeric muscle formation was observed along the anteroposterior axis of the tail, characterized by regularly spaced bar-shaped groups of mCherry^+^ cells (Fig. 3H-I). By 37.5 hpa, mAG^+^ cells at the amputation plane started accumulating at the base of the blastema (Fig. 3I). After approximately 40 hours of imaging, cells in the blastema seemed to be dying. However, subsequent time points showed continued maturation of the tail myomere muscle and a general shift from mostly mAG cells in the tail to mCherry cells.

To track migrating and dividing mAG^+^ cells, we used the Fiji plugin TrackMate (Tinevez et al., 2017). With this, we tracked the position of mAG^+^ cells 10 hours before and after each frame (Fig. 3J-M, Movie 4). This allowed us to visualize the path cells took to contribute to the regenerated tail. Interestingly, during the early blastema formation phase of the movie we observed a general trend for dorsal tail cells to migrate dorsally, ventral cells to migrate ventrally, and cells in the midline to migrate in the direction of the blastema (Fig. J-K). At later timepoints in the movie, we observed that the intense mAG^+^ cells adjacent to the amputation plane were migrating into the tail blastema (Fig. 3I), further demonstrating the local origin of blastema cells.

We quantified changes in fluorescence in mAG by dividing the regenerating tail into rectangles with a 30 μm width anterior and posterior to the amputation plane (Fig. S3). The raw integrated density for each channel was measured for each box and normalized to the total tail area within each rectangle, providing a measure of intensity per area. These results showed an increase in the intensity of mAG fluorescence in rectangles starting 30 μm anterior to the amputation plane and continuing into the regenerating tail tip, indicating that proliferation is highest in the blastema and in cells 30 μm anterior to from the blastema (Fig. 3N). Anterior to the amputation plane the mAG intensity was highest at earliest time points and steadily decreased after 60 hours of imaging (Fig. 3N). The opposite trend was observed for mCherry fluorescence, where the intensity increased after 60 hours of imaging (Fig. 3O). These results indicate an increase in the total number of cells at resting state, which may represent the termination of the rapid proliferation program employed during embryonic development.

### Multimodal imaging provides cell type identity to FUCCI tissues

A limitation to FUCCI sensors is absence of cell type information as a result of using two fluorescent proteins. This limits robust cell characterization using imaging modalities including immunohistochemistry (IHC) and fluorescence *in-situ* hybridization (FISH) along with FUCCI imaging. To overcome this limitation, we observed that mAG and mCherry can be sufficiently photobleached after imaging to allow for multimodal imaging (Fig. 4A). We first performed version 3 hybridization chain reaction FISH (V3.HCR-FISH) (Choi et al., 2018) for *Shh* using Alexa-fluor 647 on an EdU pulsed, homeostatic FUCCI spinal cord (Fig. 4B). We next photobleached the endogenous FUCCI signal and wiped the *Shh* probes with 80% formamide (Fig. 4C). Then a subsequent round of V3.HCR-FISH was performed for *Pax7* and *B3Tub* (Fig. 4D). Imaging and subsequent wiping of these probes was followed by EdU labeling (Fig. 4E). EdU signal was then removed with DNase, and IHC was performed for B3TUB (Fig. 4F). The images from each round were aligned to the DAPI image from the first round allowing imaging of four modalities (transgenic reporter, FISH, click-chemistry, and IHC) in the same tissue section (Fig 4G-J). This analysis shows that cell type identification can be performed along with the study of cell division in FUCCI tissue.

**Figure 4:**
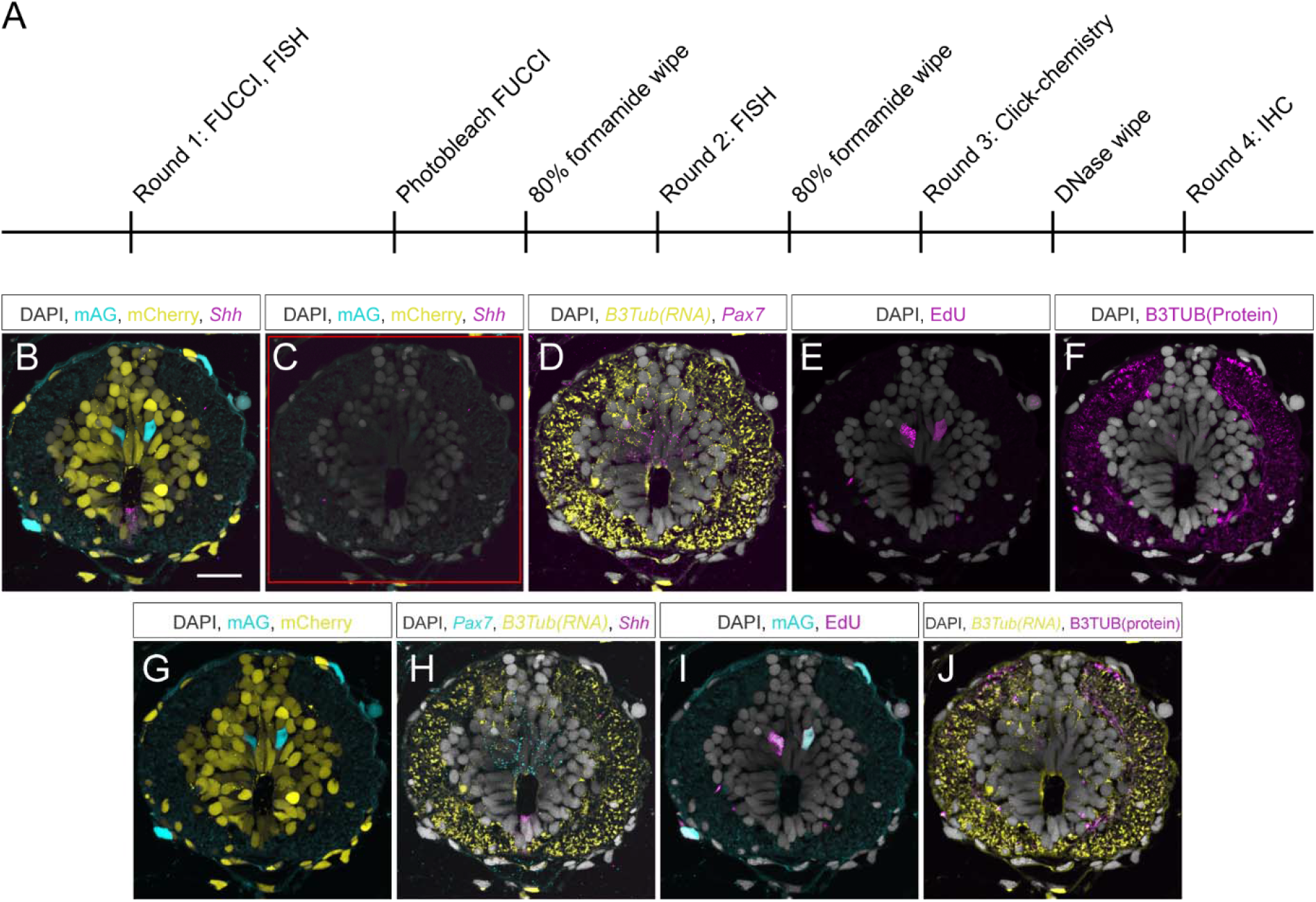
Multimodal imaging of FUCCI tissue for cell type characterization and identification of cycling cells. (A) Schematic of the staining timeline used for multimodal imaging in a homeostatic spinal cord. (B) Round one of imaging for endogenous FUCCI signal and *Shh* transcript with V3.HCR-FISH. Scale bar= 50 μm. (C) Round one of imaging after photobleaching. Red square in image represents the area photobleached. (D) Round two of imaging for *Pax7* and *B3Tub* transcript with V3.HCR-FISH. Intense signal in the white matter is autofluorescence. (E) Round three of imaging for EdU labeled cells with click-chemistry. (F) Round four of imaging for B3TUB protein with IHC. (G) Endogenous FUCCI signal in the spinal cord. (H) *Pax7*, *B3Tub* transcript, and *Shh* V3.HCR-FISH signal from rounds one and two. (I) mAG expression and EdU labeling from rounds one and three. (J) *B3Tub* transcript and B3TUB protein from rounds two and four. DAPI image used for panels B-J was obtained in round one.

### Regenerating FUCCI limbs reveal distinct regions of proliferative and non-proliferative zones in the limb blastema

To visualize cell cycling following limb amputation, we imaged uninjured limbs (Fig. S5A-B) and regenerating limbs from five animals at 1, 3, 5, 7, 10, and 14 dpa (Fig. 5A-L). To quantify the location of proliferation following amputation, we calculated the average distance between the amputation plane and the distal mCherry^+^ muscle boundary line. We found that mAG^+^ cells were abundant proximal to the amputation plane as early as 1 dpa, and that these mAG^+^ cells were located on average 243.85 μm proximal to the amputation plane (Fig. 5M). This distance from the amputation plane was significantly larger than the same measurement at 5, 7, and 14 dpa (One-way ANOVA with a Tukey-Kramer multiple comparison, p< 0.05). These findings correspond well with previous irradiation studies, where it was determined that cells from at least 500 μm proximal to the amputation plane are necessary and sufficient for regeneration (Butler, 1933). As the limb regenerates, we observed fewer mAG^+^ cells proximal to the amputation as more mAG^+^ cells accumulated within the blastema (Fig. 5M). This observation suggests that the supply of proliferating cells becomes less dependent on cells proximal to the amputation plane at later time points during regeneration.

**Figure 5:**
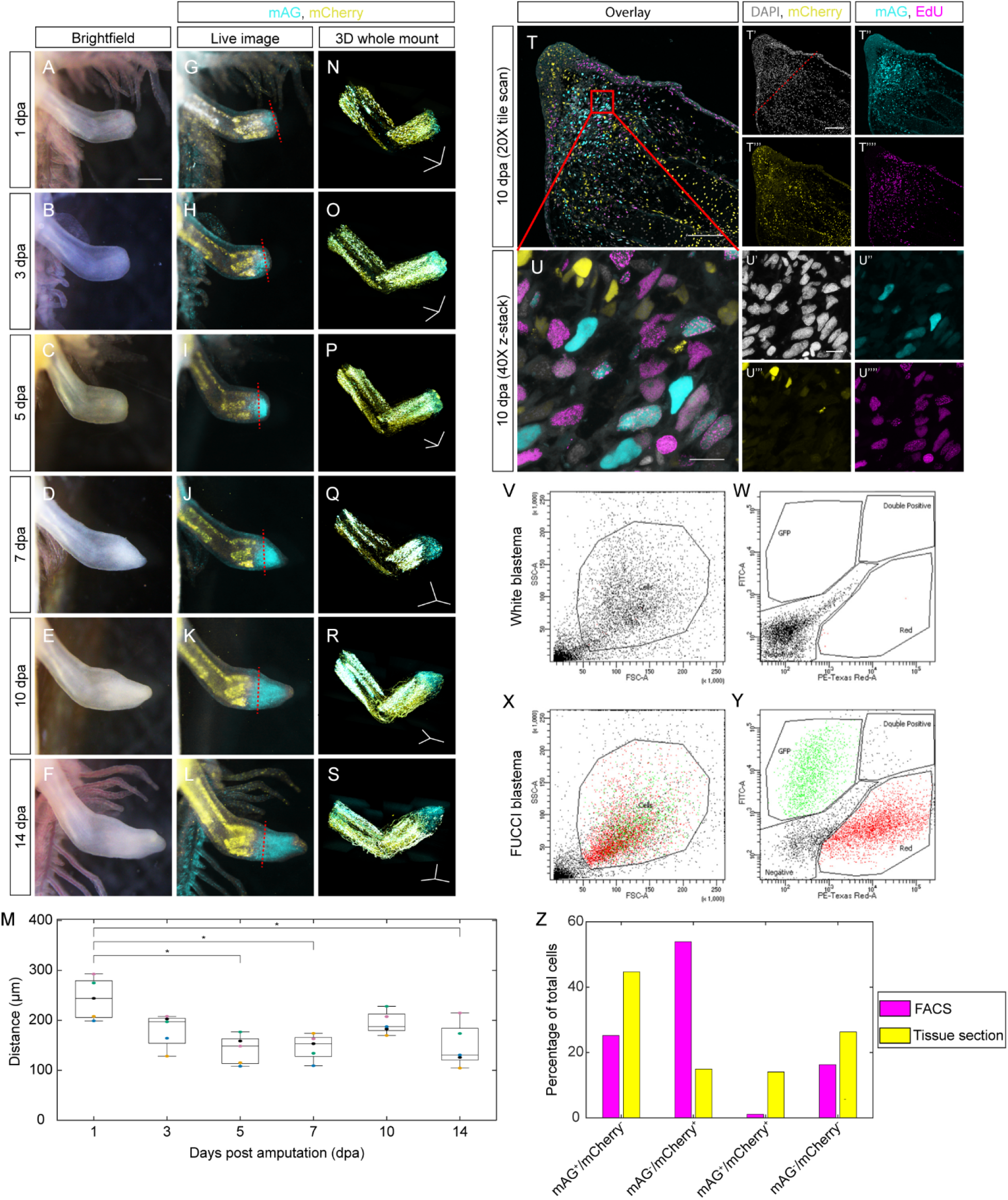
FUCCI visualization and quantification during limb regeneration. (A-F) Brightfield image of a regenerating FUCCI limb amputated through the wrist at 1, 3, 5, 7, 10, and 14 dpa. Scale bar= 0.5 mm. (G-L) mAG and mCherry fluorescence of limbs from panels A-F. (M) Quantification of the distance mAG fluorescence is observed from the amputation plane to the mCherry^+^ muscle line at each time point. Each dot color represents a replicate from a different animal. *= p value less than 0.05. (N-S) 3D, whole mount image of FUCCI limbs taken with light sheet fluorescence microscopy. Scale bars= 600 μm in each axis. (T-T’’’’) 20X tile scan of a 10 dpa FUCCI blastema pulsed with EdU for three hours. Scale bars= 150 μm. (U-U’’’’) 40X z-stack of the EdU pulsed blastema mesenchyme. Scale bars= 25 μm. (V-Y) Flow cytometry analysis of blastema cells from white strain (V-W) and FUCCI (X-Y) axolotls. A forward scatter (FSC-A) and side scatter (SSC-A) plot was used to gate for the cell population (V, X). Stage of cell cycle was determined using a PE-Texas Red versus FITC scatter plot displaying only the gated cell population (W, Y). (Z) Quantification of mAG^+^/mCherry^−^, mAG^−^/mCherry^+^, mAG^+^/mCherry^+^, and mAG^−^/mCherry^−^ cell populations in FACS sorted blastemas and tissue sections.

To determine the cell type of the mAG^+^ and mCherry^+^ cells in regenerating limbs, we visualized FUCCI probe expression in whole mount with light sheet fluorescence microscopy (Fig. 5N-S, Fig. S5 C-L, Movies 5-6). The majority of uninjured tissue including fibroblasts, epithelial cells, and chondrocytes were mCherry^+^. Most muscle cells observed were mAG^+^/mCherry^+^, suggesting cell cycle arrest at the restriction point (R-point) of the cell cycle. This finding is consistent with a similar G1/S arrest in FUCCI mouse cardiomyocytes (Alvarez et al., 2019). Very few mAG^+^ cells were observed in uninjured tissue (Fig. S5B-C,E). At 1 dpa, we observed several mAG^+^ cell types, including the wound epithelium, perichondrium, and in some fibroblasts of the mesenchyme (Fig. S5F). These cell types appear to remain mAG^+^ until blastema formation at 7 dpa, where fewer chondrocytes, perichondrial cells, and epithelial cells are mAG^+^ (Fig. SG-I). From 7-14 dpa, most of the mAG^+^ cells are located within the mesenchyme, further showing that cells proximal to the amputation plane less frequently proliferate at later time points during limb regeneration (Fig. SJ-K). At 10 and 14 dpa, we observed a small population of cells at the distal-most tip of the blastema that were mAG^−^/mCherry^+^ (Fig. 5K-L). We sectioned EdU pulsed 10 dpa blastemas for histological analysis (n=3) and found that this mAG^−^/mCherry^+^ population was composed of the distal-most epithelial cells of the AEC (Fig. 5T). In one sample, a small number of these cells was observed in the distal-most portion of the blastema mesenchyme (Fig. 5T). We then sectioned 14 dpa blastemas (n=3) and observed the presence of these distal-most, mesenchymal mAG^−^/mCherry^+^ cells in all three samples (Fig. S6), suggesting that this population of cells is more abundant at later stages during regeneration. The observation that the highest proliferation levels being in the middle-proximodistal region of the blastema has been observed by others in salamanders (Farkas et al., 2016; McCusker et al., 2015b) and in regenerating zebrafish fins (Hirose et al., 2014).

For quantification of blastema cells, we performed flow cytometry analysis on 10 dpa FUCCI blastemas (n=10) (Fig. 5V-Y). In total, 5,682 cells were analyzed of the total 10,000 events. Of these cells, 25.2% mAG^+^/mCherry^−^, 53.9% were mAG^−^/mCherry^+^, 1.1% were mAG^+^/mCherry^+^, and 16.3% were mAG^−^/mCherry^−^. Although, these results do not exactly correspond with our tissue section quantification (n=3), where we found that 44.68% mAG^+^/mCherry^−^, 14.96% were mAG^−^/mCherry^+^, 14.06% were mAG^+^/mCherry^+^, and 26.3% were mAG^−^/mCherry^−^, it is not surprising considering some mCherry^+^ cells just proximal to the blastema were collected for dissociation (Fig. 5Z). Additionally, the amount of autofluorescent blood as well as the cessation of proliferation that likely occurs as the cells are dissociated prior to FACS analysis may contribute to the discrepancy.

### Spinal cord amputation induces a proliferation response 5 mm from the injury

To determine the location of proliferating cells along the anteroposterior (AP) axis of the regenerating spinal cord, we collected EdU pulsed FUCCI tissue sections (n=4) at various locations along the AP axis with respect to the most posterior tip of the regenerated cartilaginous rod (Fig. 6A). The amputation plane is located between the 250 μm anterior and 500 μm anterior sections, as the notochord was identified 500 μm anterior to the cartilaginous rod but not at the 250 μm anterior section. For comparison, we sectioned spinal cords from non-regenerating, homeostatic FUCCI animals (n=5). Quantification included cells within the boundary of the meninges that surrounds the spinal cord but excluded the meningeal cells themselves (Fig. 6B). mAG^+^/mCherry^−^ cells were consistently around 40% of total cells posteriorly and progressively declined anteriorly (Fig. 6C-L), which may indicate that cell proliferation is most abundant at or posterior to the cartilaginous rod tip. This is accompanied by an increase in the number of mAG^−^/mCherry^+^ cells anteriorly along the regenerating AP axis (Fig. 6C-L), suggesting a shortened G1 phase in regenerating cells which is supported by previous studies (Rodrigo Albors et al., 2015). Furthermore, we observed a significant increase in the number of mAG^+^/mCherry^−^ cells located 5000 μm from the regenerated cartilaginous rod compared to uninjured spinal cords (Two tailed Student’s t-Test assuming unequal variances, p=0.0043), suggesting that spinal cord injury induces an increase in cell cycling beyond 500 μm anterior to the amputation plane (Fig. 6L). Our results also indicate that the relative abundance of cells in S or G2/M as indicated by mAG^+^/EdU^+^ or mAG^+^/EdU^−^, respectively, is unchanged across the AP axis (Fig. 6M), suggesting that the ratio of S:G2 does not significantly change across the regenerating AP axis. However, a significant difference is detected between the total number of mAG^+^/EdU^+^ cells detected in sections 5000 μm from the cartilaginous rod tip and uninjured spinal cords (Two tailed Student’s t-Test assuming unequal variances, p= 0.001) (Fig. 6M). A statically significant difference between the number of mAG^+^/EdU^−^ cells at these regions was not detected (Two tailed Student’s t-Test assuming unequal variances, p= 0.143) (Fig. 6M). Taken together, our results indicate that spinal cord injury induces an increase in the number of cycling cells along the AP axis 5 mm from the injury compared to uninjured controls.

**Figure 6:**
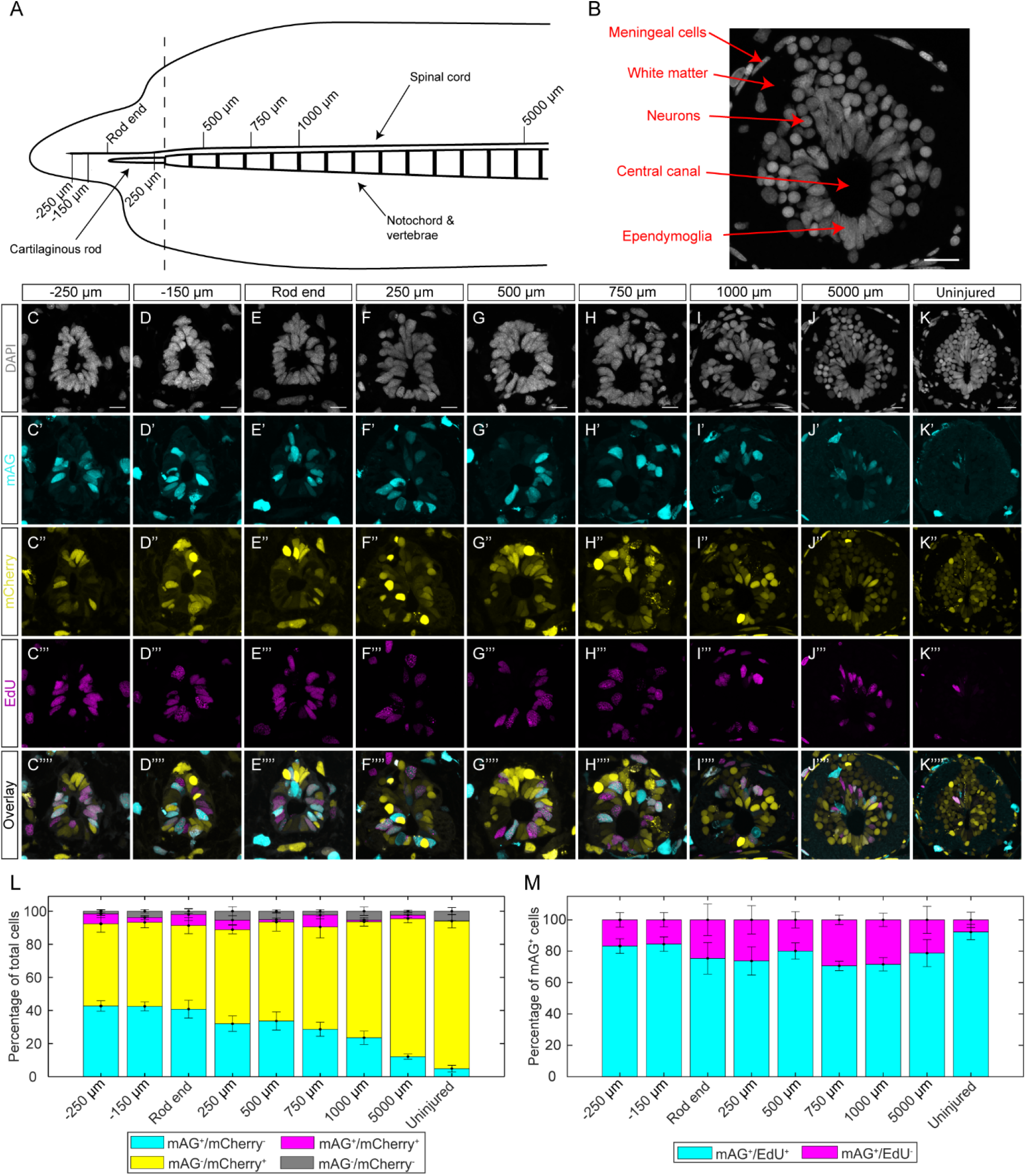
Spinal cord amputation induces a proliferative response 5 mm from the amputation plane. (A) Schematic of the experiment. (B) Cell types of the spinal cord. Scale bar= 25 μm. (C-K) Individual channels for spinal cord cross sections pulsed with EdU. Scale bars for panels C-J= 25 μm. Scale bar for panel K= 50 μm. (L) Total cell quantification across the regenerating AP axis. (M) mAG^+^ cell characterization across the regenerating AP axis.

## Discussion

Many fundamental questions remain unanswered regarding cell proliferation during appendage regeneration. How cell cycle dynamics change during regeneration compared to uninjured limbs, if the cell cycle length is unique to individual regenerating organs, and if the cell cycle is regulated differently during development versus regeneration are among some of the many outstanding questions. Previously, these questions were addressed using a combination of thymidine analogs like EdU and BrdU, pHH3 antibody labeling, and mitotic figures. However, none of these methods provide information about multiple cell cycle stages and are unable to be used for live imaging. FUCCI axolotls provide a powerful means to address these questions in detail while improving the existing toolbox for the study of cell cycle dynamics during tissue regeneration. Furthermore, a major advantage of deploying FUCCI sensors in the axolotl is the amenability of axolotl embryonic tissue to grafting, which has been shown to be successful for limb connective tissue, muscle cells, epithelium, Schwann cells, vasculature, neural stem cells, neural crest cells, and teeth primordium (Epperlein et al., 2012; Kragl et al., 2009; Kragl and Tanaka, 2009; Nacu et al., 2013). Grafting CAG FUCCI tissue onto white embryos will allow for tissue-specific expression without the need for the generation of new transgenic animals with a tissue specific promoter driving FUCCI expression.

While FUCCI sensors in the axolotl are highly useful, some limitations exist. One issue of our transgenic line is the variable expression across animals and across tissues. This is particularly obvious in adult animals, where some animals seem to strongly express only one fluorescent protein. Continuous breading of the line to d/d mates should decrease variability across siblings. The use of EdU in FUCCI tissue also has small limitations. First, the number of mAG^+^/mCherry^−^/EdU^−^ cells may be underrepresented and the number of mAG^+^/mCherry^−^/EdU^+^ may be overrepresented; after a three hour pulse of EdU, cells labeled in late S phase will transition to G2 phase prior to collection. The severity of this issue can be reduced by collecting tissue sooner than 3 hours after the EdU pulse, but this problem is theoretically always possible. Second, we observed the presence of mAG^−^/mCherry^+^/EdU^+^ and mAG^−^/mCherry^−^/EdU^+^ cell populations in our samples. Our quantification suggests that these populations are not highly abundant, and we speculate that they are detected as a result of the cells not expressing the FUCCI construct or DNA damage, as EdU is known to be incorporated into cells undergoing DNA repair (Verbruggen et al., 2014).

In our study, we highlight the versatility of imaging FUCCI tissue with live imaging, whole mount imaging, and multimodal imaging. To our knowledge, we present the first real time, *in-vivo* movie of blastema formation in regenerating axolotl tissue. The approaches used in this study will be helpful for other groups trying to live image appendage regeneration in real time, but methods should be optimized. As the blastema was growing, we observed a large number of dying cells and cells sluffing from the blastema. After removing the larvae from the agarose, we also noticed that the blastema was misshapen. We do not think this is an accurate representation of tail blastema growth and is more likely a result of the blastema growing in the agarose. We also visualize FUCCI expression in whole mount regenerating limbs with light sheet fluorescence imaging. We attempted to visualize these limbs in 3D after staining for EdU, but the EdU signal from the 647 channel was too strong and bled into the mCherry signal. With better filtering, we expect to be able to perform 3D multiscale analysis of macromolecule synthesis in FUCCI tissue. Finally, we outline a method for cell characterization in transgenic tissue with multimodal imaging. These proof of concept experiments demonstrate the amount of cell type information one can acquire from a single FUCCI tissue section. We foresee better cell type characterization via multimodal imaging with more rounds of FISH, multiple macromolecule analogs with unique click-it compatible functional groups (Duerr et al., 2020), and multiple primary antibodies raised in different species. One potential limitation of this method is the inefficiency of photobleaching FUCCI signal in large tissues. The spinal cord is an ideal organ for this analysis, as it can easily fit into a single 20X frame. Thus, photobleaching is contained to one single tile. If using larger tissue like a limb blastema, many more tiles may require photobleaching and potentially at a lower magnification, both of which will increase the time necessary to completely photobleach the FUCCI signal. Overall, our method can be used for robust characterization of cycling cells during tissue regeneration.

## Methods

### Animal procedures

All transgenic animals were bred at Northeastern University, and all procedures and surgeries were approved by the Northeastern University Institutional Animal Care and Use Committee. Surgeries were performed while axolotls were anaesthetized in 0.01% benzocaine. EdU was administered via intraperitoneal injection at 8.0 ng/g animal and samples were collected three hours after injection.

### Transgenesis

Transgenesis was performed via I-SceI meganuclease digestion according to Khattak 2009. Briefly, 1 μg of purified, CAG FUCCI plasmid was mixed in solution with 2 μL NEB Cutsmart buffer and1 μL I-SceI enzyme, filled to a final volume of 10 μL with nuclease free water to generate the FUCCI injection cocktail. Single cell, d/d axolotl embryos were injected with 5 nL of FUCCI injection cocktail and grown to stage 45 for phenotype assessment. Due to the beta-actin promoter in CAG, the most intense FUCCI expression was observed in myomeric muscle of developing tails. Larvae with strong, ubiquitous expression were identified and grown to sexual maturity.

### Histology and staining

Samples were fixed in 4% PFA overnight at 4°C, and after washing with 1X PBS three times for 5 minutes, samples were cryoprotected in 30% sucrose until equilibrated. Samples were then placed in OCT and frozen at −80°C. Frozen samples were sectioned with a cryostat to obtain 10 μm sections. Slides were then baked at 65°C for 15 minutes. Residual OCT was removed from slides by placing in water for 5 minutes at room temperature. From this step, the slides are ready for click-chemistry, IHC, or FISH:

#### EdU labeling via click chemistry

For EdU detection, we used an Alexa-fluor 647 azide plus probe from https://clickchemistrytools.com (Product number: 1482). The 1 mL click-it cocktail was made as follows: 885 μL 1X Tris, 10 μL 50 mM CuSO_4_ (0.5 mM final), 2 μL Alexa-fluor 647 azide plus (2 μM final), 100 μL 100 mM sodium ascorbate (10 mM final). This cocktail was applied to slides for 30 minutes at room temperature.

#### IHC

Slides were incubated in blocking buffer (15 μL goat serum in 1 mL 1X PBS) for 30 minutes. Rabbit anti-pHH3 antibodies were diluted in blocking buffer at a 1:400 concentration and applied to slides overnight at 4°C. Slides were washed three times for 5 minutes each with 1X PBS, and 647 anti-rabbit secondary antibodies (diluted 1:500 in 1X PBS) were applied to slides for 30 minutes at room temperature.

#### Multi-round V3.HCR-FISH

All of the following steps are conducted using RNase free reagents. Slides were placed in 100% ethanol at room temperature for 1 hour. Following three 5 minute washes with 1X PBS, slides were prehybridized with hybridization buffer (Molecular Instruments) for 30 minutes at 37°C. Probe stocks for a particular transcript of interest were made to contain 1 μM of each oligo in 200 μL of RNase free water. Probe sequences for *Shh*, *B3Tub*, and *Pax7* are provided in the supplementary material. Probe stocks were further diluted 1:200 in hybridization buffer and applied to slides overnight at 37°C. Slides were washed with formamide wash buffer (Molecular Instruments) three times for 15 minutes at 37°C to remove unbound probe, then washed twice with 5X SSCT (20X saline sodium citrate with 0.1% Tween 20) for 15 minutes at room temperature. Amplification buffer (Molecular Instruments) was then applied to the slides for 30 minutes at room temperature. Fluorescent hairpins for each initiator (Molecular Instruments) were prepared by heating H1 and H2 hairpins to 95°C for 90 seconds. Hairpins were allowed to cool to room temperature in the dark, then diluted 1:50 in amplification buffer and applied to slides overnight at room temperature. Slides were then washed twice for 30 minutes with 5X SSCT at room temperature.

After these protocols, cell nuclei were stained with DAPI (2.86 μM) for five minutes at room temperature. Following a five minute 1X PBS wash at room temperature, slides were mounted with SlowFade gold antifade mountant (Thermo S36936) and imaged using a Zeiss LSM 800 confocal microscope.

### Live FUCCI imaging

Larvae used for live imaging were mounted in a 50 × 9 mm petri dish in 0.3% low melt agarose diluted in 0.005% benzocaine. All live images were acquired using a Zeiss LSM 880 confocal microscope fitted with a humidification chamber to prevent sample desiccation. Larvae were imaged at 10X magnification. For live imaging of tail regeneration, we imaged two adjacent tiles to accommodate for growth during imaging. Additionally, to accommodate for cells moving in and out of the focal plane, we imaged four planes in the z axis and merged these planes together in a maximum intensity projection. To prevent photobleaching of the FUCCI probes, we used 1.0% laser power for each channel. Larvae were removed from agarose after imaging and placed in salamander housing water. Larvae were swimming and feeding one week after imaging with no visible signs of illness or distress.

### Multimodal imaging

EdU pulsed FUCCI spinal cords were collected as outlined above. In the first round of multimodal imaging, we performed V3.HCR-FISH for *Shh* with 647 hairpins. The endogenous FUCCI signal and V3.HCR-FISH was then imaged. To photobleach the FUCCI signal, the 488 and 594 lasers were set to 100% laser power and were directed onto the spinal cord for 40 minutes. We found that the DAPI signal was weakened after this photobleaching, but still present. The V3.HCR-FISH signal was sufficiently photobleached, but to ensure *Shh* probes were not amplified in the subsequent round of FISH, the slides were washed with 80% formamide four times for 15 minutes each at 37°C. The slides were washed in 5X SSCT twice for 15 minutes each at room temperature, prehybridized with hybridization buffer for 30 minutes at 37°C, and rehybridized with *Pax7* and *B3Tub* probes for the second round of multimodal imaging. These probes were amplified with 647 and 488 hairpins, respectively. Slides were imaged, and probes were again removed with four 15 minute washes of 80% formamide at 37°C. Slides were washed three times with 1X PBS, and the click-it cocktail outlined above was used with Alexa-fluor 647 azide plus probes for EdU labeling in the third round of multimodal imaging. Slides were imaged and treated with DNase I (NEB M0303) overnight at room temperature. DNase I was applied to slides without buffer, and enough was used to cover the entire section being imaged. The next day, we performed IHC with rabbit anti-B3TUB antibodies (1:500) and applied anti-rabbit 647 antibodies the subsequent day. Slides were then imaged for the final round of multimodal imaging. Adobe Photoshop was used to align the images from each round onto the original DAPI image. All images were obtained with a Zeiss LSM 800 confocal microscope.

### Whole mount FUCCI imaging

To prepare whole mount tissue, limbs were fixed in 4% PFA overnight at 4°C. Limbs were then washed with 1X PBS three times for 5 minutes, and dehydrated in an increasing methanol series (25% MeOH/75% 1X PBS, 50% MeOH/50% 1X PBS, 75% MeOH/25% 1X PBS, each step for 5 minutes at room temperature), and stored in 100% methanol at −20°C for up to 6 months prior to imaging. Once ready to be imaged, the limbs were rehydrated in a decreasing methanol series (75% MeOH/25% 1X PBS, 50% MeOH/50% 1X PBS, 25% MeOH/75% 1X PBS, each step for 5 minutes at room temperature) and washed once with 1X PBS for 5 minutes. The samples were washed three times with 1X PBST (Triton 100X) for 5 minutes at room temperature. We found that a 90 minute 0.5% trypsin treatment at room temperature with rocking improved light penetration of FUCCI samples without appreciable changes in intensity of mAG and mCherry. After trypsin treatment, samples were washed with deionized water three times for 5 minutes at room temperature. The limbs were then placed in 100% acetone for 20 minutes at −20°C. Afterwards, the samples were incubated in deionized water for 10 minutes at room temperature. Samples were again washed with 1X PBS three times for 5 minutes, mounted in 1.5% low melt agarose, and refractive index matched with EasyIndex RI 1.465 (LifeCanvas Technologies) overnight at 4°C. Three dimensional images were obtained using a Zeiss light sheet Z1 microscope with Zen software. All post processing for visualization was performed using Arivis software.

### Cell dissociation and Flow Cytometry

Ten 10 dpa blastemas from white strain or FUCCI animals (3-5cm snout to tail tip), were collected and pooled together in a 6-well plate. The wound epithelium was not surgically removed. Blastemas were incubated on ice in 0.35 mg/mL Liberase for 20 minutes during transfer to FACS core facility and were then incubated at room temperature with gentle agitation. Every 10 minutes, the tissue was manually dissociated with forceps by teasing the tissue apart. This was repeated until there was a sufficient single cell suspension (checked under the microscope) while the wound epithelium remained intact. Using 1 mL of 80% PBS, the cell suspension was filtered using a 35 μm filter tube, leaving behind the wound epithelium. The strainer was washed with an additional 1 mL of 80% PBS.

The filtered single cell suspensions were run on a BD FACSAria Fusion Cell Sorter (UMass Boston Flow Cytometry Core) using the 100 μm nozzle and the FSC 2.0 ND filter. The gates for the blastema cell population were set on the blastema cells from the white animal using forward and side scatter. Using the forward scatter and side scatter plot, the cell population was gated to separating it from debris and doublets. A sample of the gated population was sorted, and the presence of singlet cells was conformed with microscopy. The gated cell population was then analyzed on a PE-Texas Red and FITC scatter plot to gate the fluorescent negative cell population. The same gates were then used when the FUCCI blastema cells were filtered and run through the cell sorter. Gates were added to quantify the red, green, and double positive populations. Gates for fluorescent populations were also confirmed by sorting and validating cell populations with fluorescent microscopy.

### Data analysis

FUCCI^+^ cells were quantified either manually in Adobe Photoshop or with Cellpose (Stringer et al., 2021) combined with custom Fiji scripts (Schindelin et al., 2012), which are available in the supplementary information. Fiji scripts for quantification of mAG/mCherry intensity changes during tail regeneration are also available in the supplementary information. Limb blastema amputation plane to mCherry^+^ muscle line measurements were made using the InteredgeDistance macro on Fiji. Data processing and statistical analysis were conducted on Microsoft Excel and Matlab.

## Acknowledgements

The authors would like to thank Jackson Griffiths for assisting with transgenic animal care during the COVID-19 pandemic lockdown, Ester Comellas for assistance with data analysis, both Guoxin Rong and Alex Lovely for imaging assistance, and Malcom Maden for early conceptual discussions on the project. Images were obtained from the Northeastern University Chemical Imaging of Living Systems core. We thank the Institute for Chemical Imaging of Living Systems at Northeastern University for consultation and imaging support. Non-transgenic animals were obtained from the Ambystoma Genetic Stock Center funded through NIH grant P40-OD019794.

---

**Figure S1:**
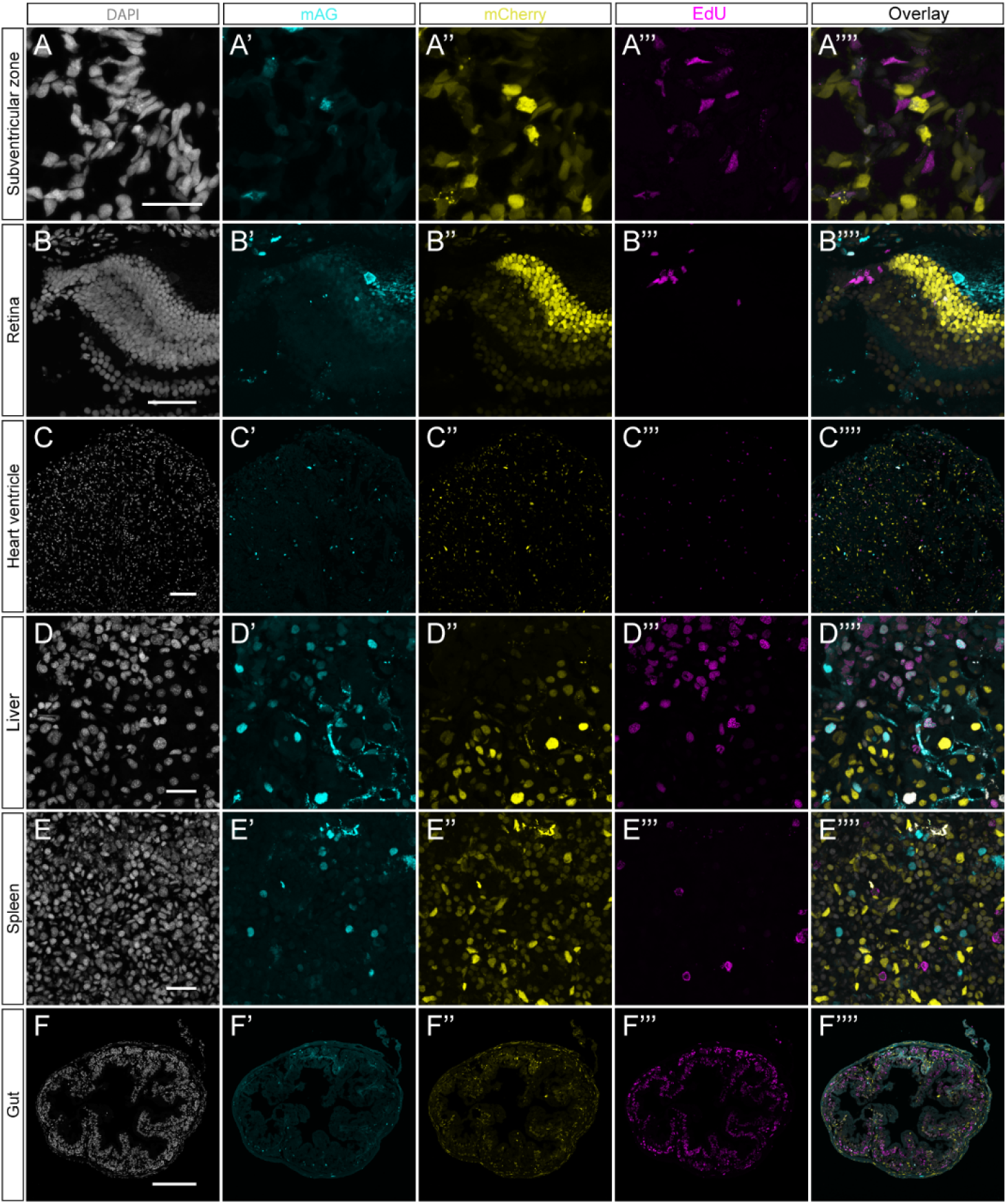
Single channel images from Figure 1. (A-F’’’’) Individual channels from panels I-J in Fig. 1 with EdU staining. Scale bars are identical as in Fig. 1.

**Figure S2:**
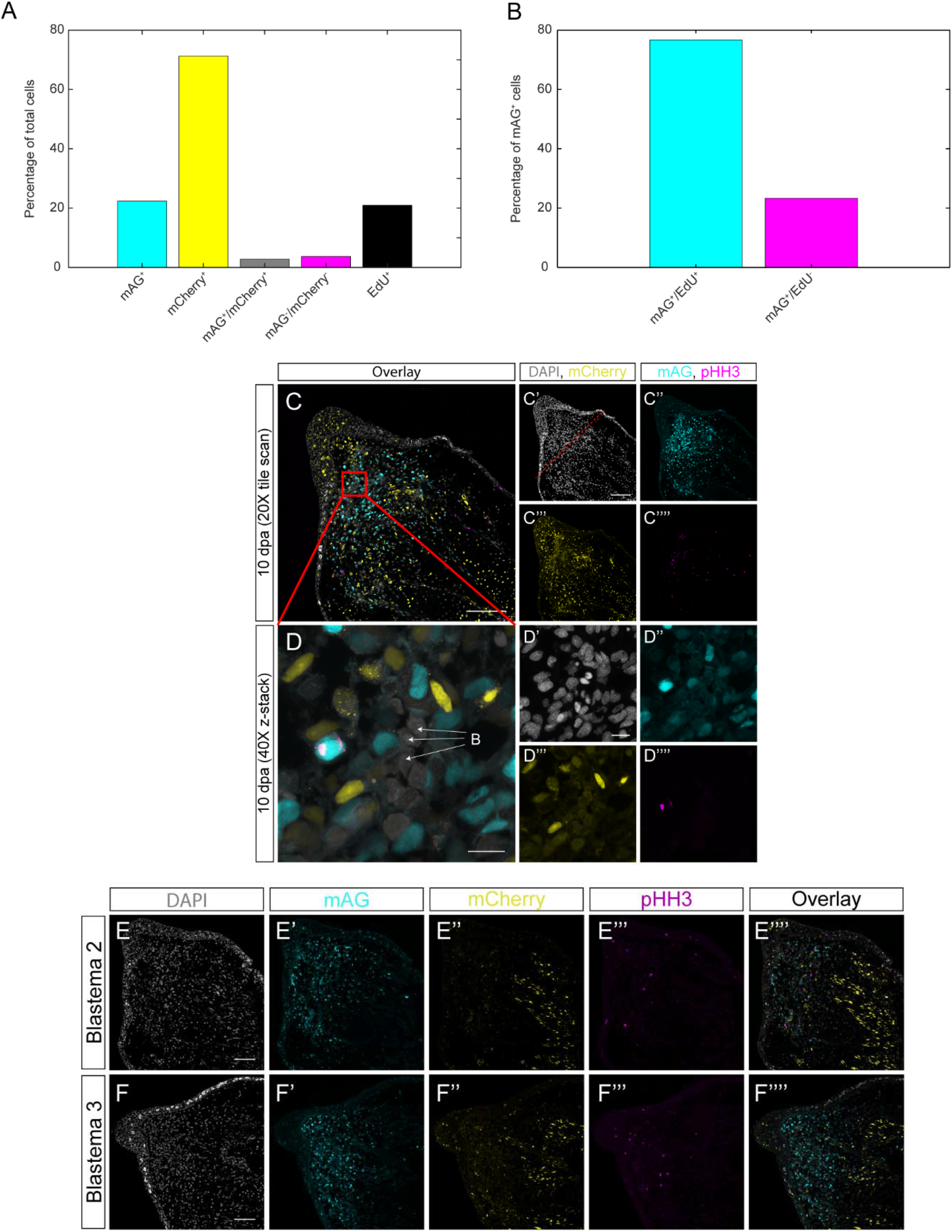
Additional information from FUCCI validation. (A) Total cell characterization of the 2547 cells from the EdU pulsed 14 dpa regenerating spinal cords. (B) Quantification of the number of mAG^+^/EdU^+^ and mAG^+^/EdU^−^ cells from the EdU pulsed 14 dpa regenerating spinal cords. (C-C’’’’) 20X tile scan of a 10 dpa FUCCI blastema stained with pHH3. Scale bars= 150 μm. (D-D’’’’) 40X z-stack of pHH3 stained blastema mesenchyme. B= blood cells. Scale bars= 25 μm. (E-F) Two additional replicates of 10 dpa FUCCI limb blastemas stained for pHH3. Scale bars= 100 μm.

**Figure S3:**
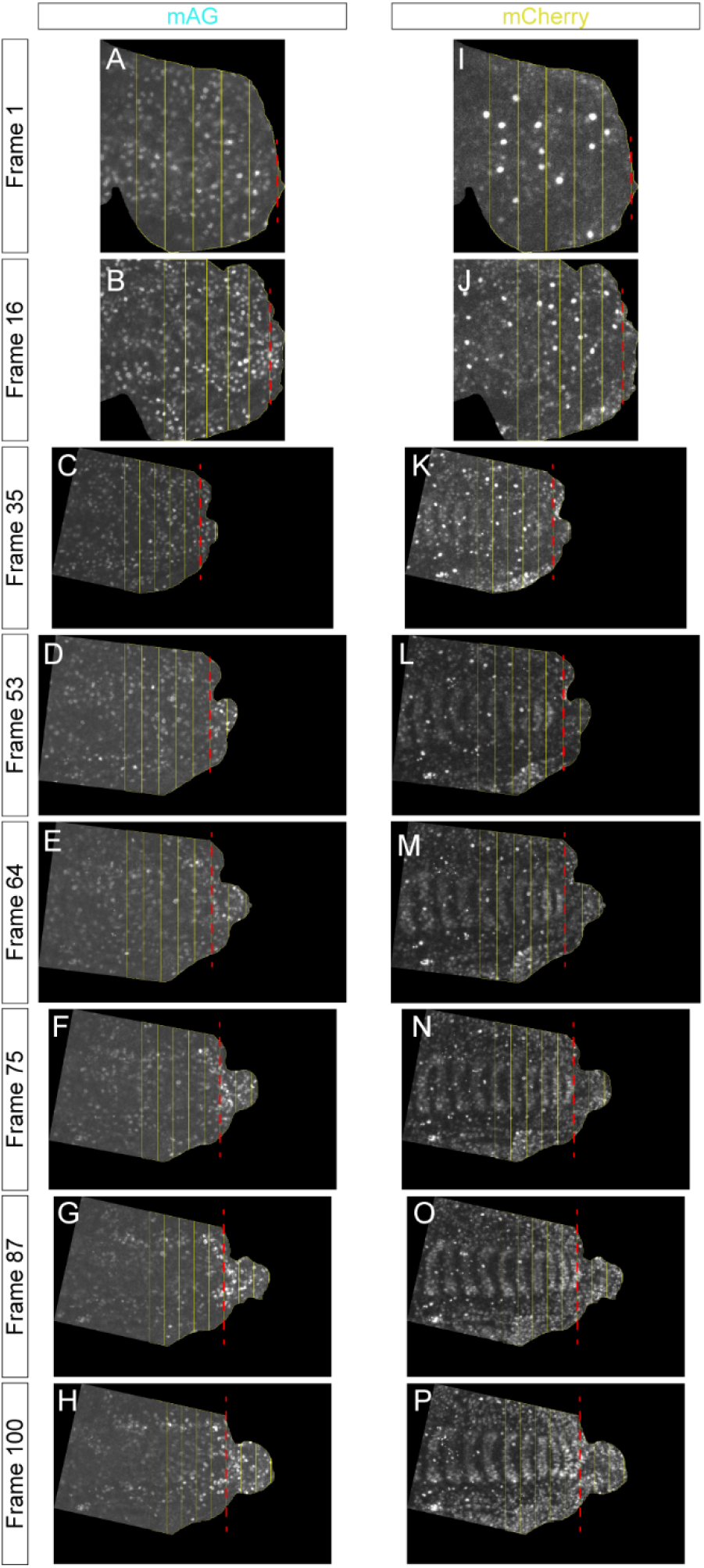
Tail regeneration live image quantification boxes. (A-H) Frames with quantification boxes for mAG. (I-P) Frames with quantification boxes for mCherry. Red dashed line indicates amputation plane on each frame. Raw integrated density was mesaured for each box and divided by total box area.

**Figure S4:**
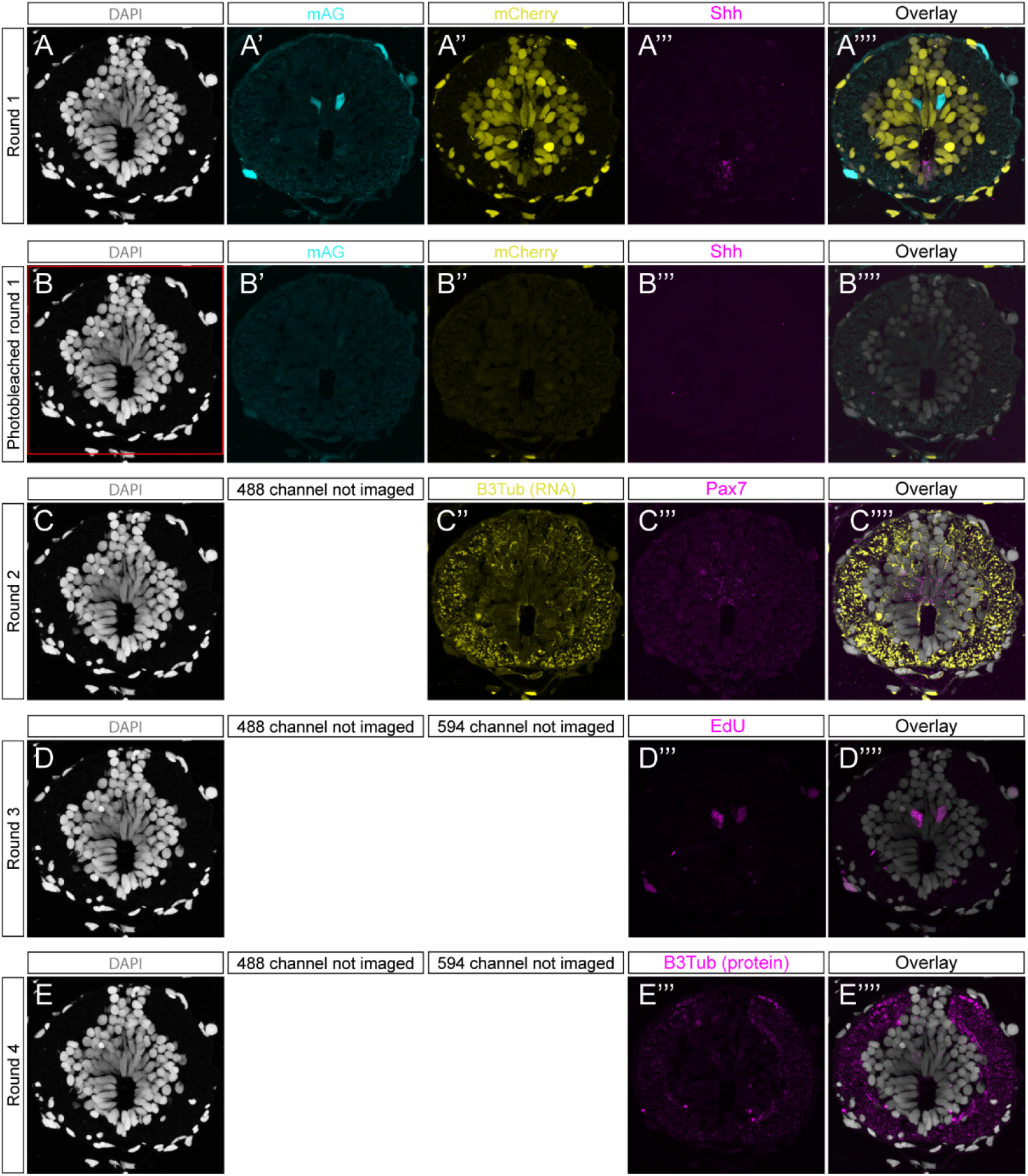
Single channel images from Figure 4. (A-E’’’’) Individual channels from each round of multimodal imaging.

**Figure S5:**
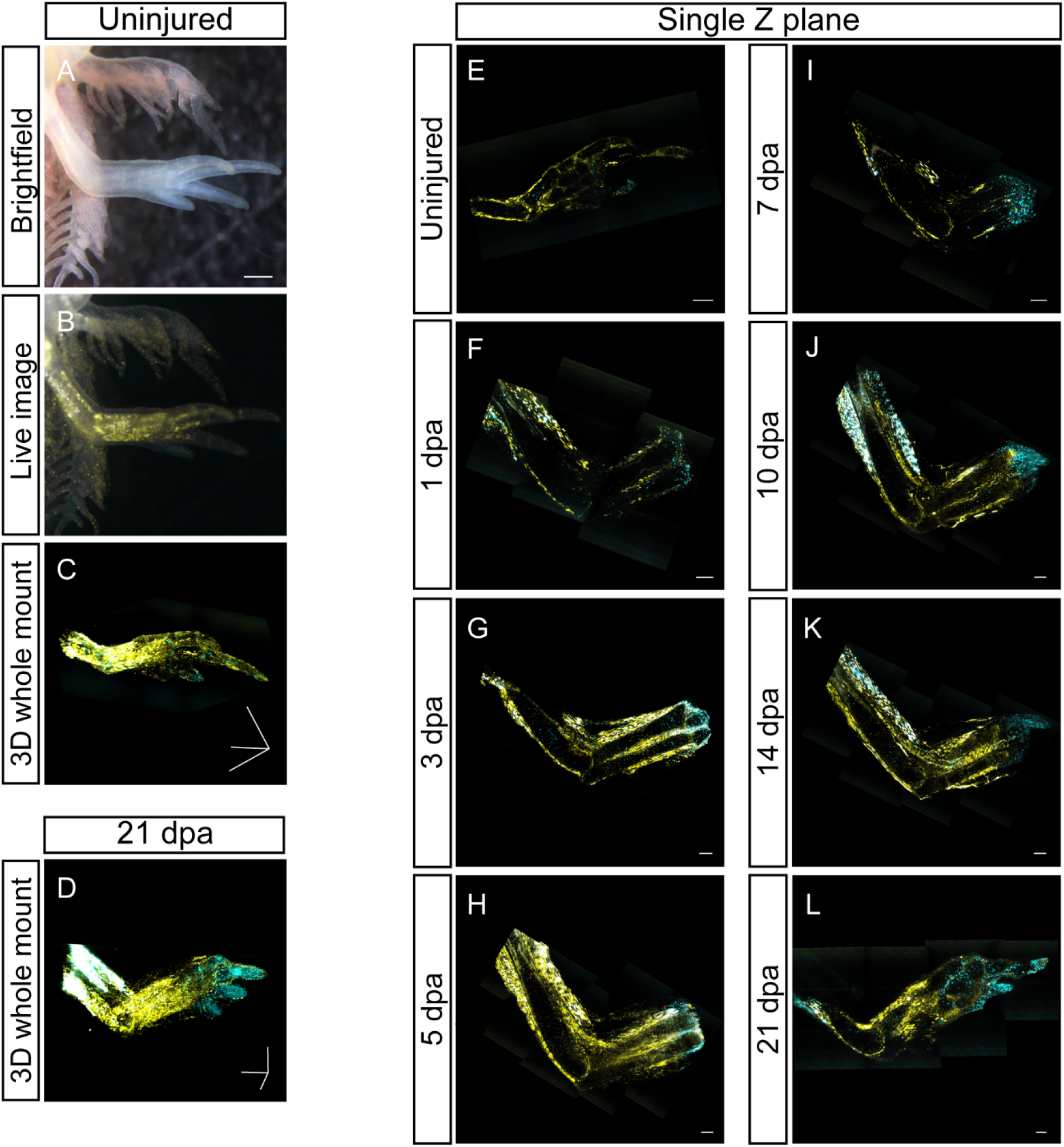
Additional FUCCI limb regeneration information. (A) Brightfield image of an uninjured FUCCI limb. Scale bar= 0.5 mm. (B) mAG and mCherry fluorescence of limb from panel A. (C-D) 3D, whole mount image of uninjured (C) and 21 dpa (D) FUCCI limbs taken with light sheet fluorescence microscopy. Scale bars= 600 μm in each axis. (E-L) 2D z slices of whole mount images from panels C-D and Figure 5 panels N-S. Scale bars= 200 μm.

**Figure S6:**
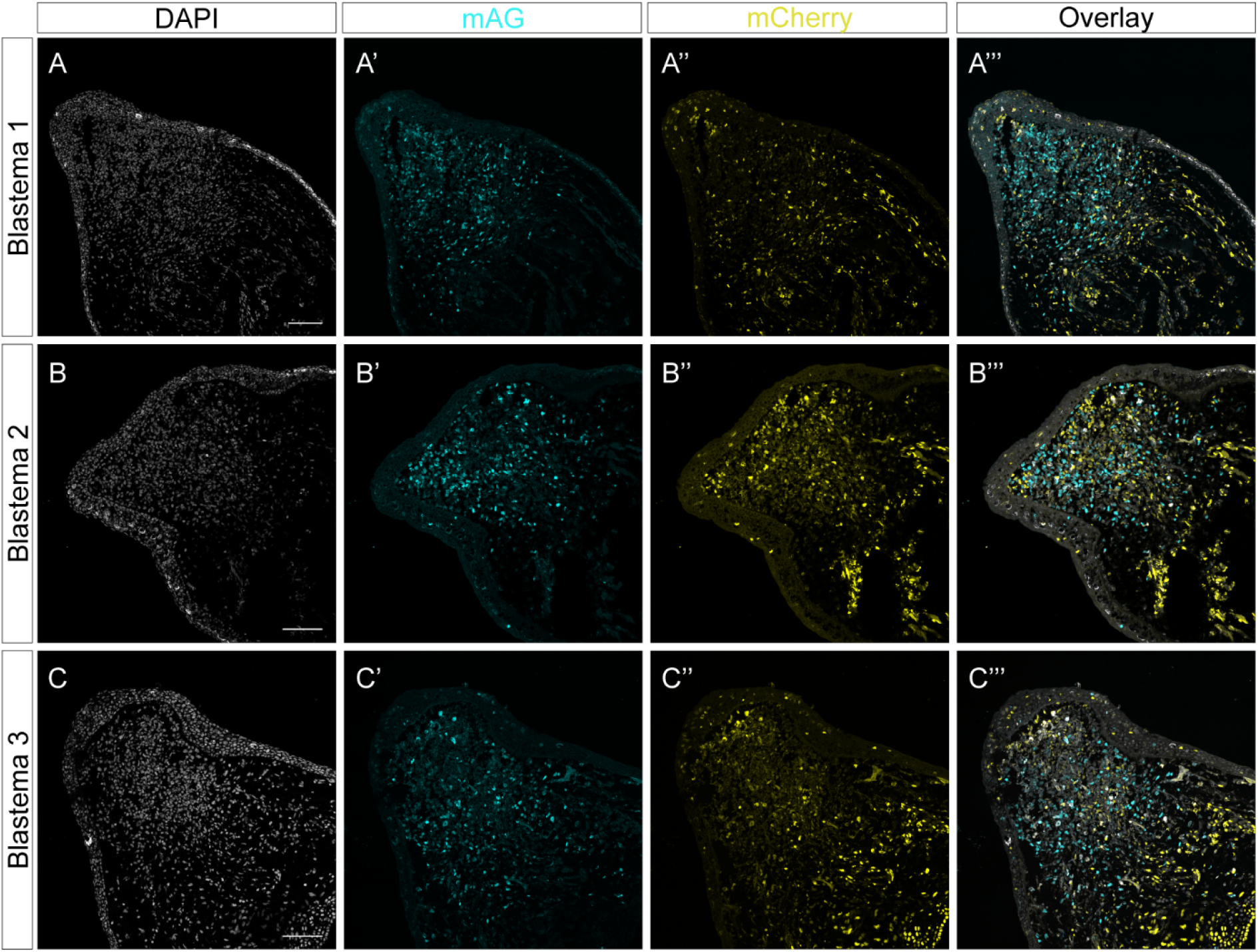
14 dpa regenerating FUCCI limbs. (A-C’’’) Single color channels for three replicates of 14 dpa FUCCI limbs. Scale bars= 100 μm.

**Movie 1: 16 hour live image of dividing epithelial cells in a stage 32 FUCCI larva**

**Movie 2: Dividing mAG^+^ epithelial cell from a stage 32 larva**

**Movie 3: 60 hour live image of tail regeneration from a stage 36 FUCCI larva**

**Movie 4: mAG tracks for each frame from Movie 3**

**Movie 5: Whole mount uninjured FUCCI limb blastema**

**Movie 6: Whole mount 21 dpa regenerating FUCCI limb blastema**

